# A simplified and highly efficient cell-free protein synthesis system for prokaryotes

**DOI:** 10.1101/2025.09.12.675764

**Authors:** Xianshengjie Lang, Changbin Zhang, Jingxuan Lin, Zhe Zhang, Wenfei Li

## Abstract

Cell-free protein synthesis (CFPS) systems are a powerful platform with immense potential in fundamental research, biotechnology, and synthetic biology. Conventional prokaryotic CFPS systems, particularly those derived from *Escherichia coli* (*E. coli*), often rely on complex reaction buffers containing up to thirty-five components, limiting their widespread adoption and systematic optimization. Here, we present an optimized *E. coli* cell-free protein synthesis (*e*CFPS) system, which is significantly streamlined for high efficiency. Through systematic screening, we successfully reduced the essential core reaction components from 35 to a core set of 7. The thorough optimization of these seven key components ensured that protein expression levels were not only maintained but even substantially improved. Furthermore, we developed a much simpler procedure for preparing the bacterial cytosolic extracts, a “fast lysate” protocol that eliminates the traditional time-consuming runoff and dialysis steps, thereby enhancing the overall accessibility and robustness of *e*CFPS. This optimized and user-friendly *e*CFPS efficiently synthesizes challenging proteins, including functional, self-assembling vimentin, and active restriction endonuclease *Bsa*I despite its strong cytotoxicity, and serves as a powerful tool that will facilitate diverse applications in basic life science research and beyond.

## Introduction

Cell-free protein synthesis (CFPS) offers a powerful and flexible platform for biological research and biotechnological applications by reproducing the cellular protein synthesis process in an *in vitro* environment^1,2^. Compared to traditional cell-based expression systems, CFPS possesses several notable advantages, including rapid reaction kinetics, ease of operation, independence from cell culture, high tolerance to toxic proteins, and facile incorporation of non-canonical amino acids^3–5^. These characteristics position CFPS for broad applications in protein engineering, high-throughput screening, synthetic biology, and diagnostic reagent development^6,7^.

Prokaryotic CFPS systems, especially those based on *E. coli* lysates, have garnered significant attention for their high protein synthesis yields and compatibility with genetic manipulations^8–13^. They have played a pivotal role in early molecular biology research and continue to be relevant in industrial protein production and biosensor development^1,14–16^. However, for a long time, most prokaryotic CFPS protocols have relied on complex buffer systems containing a large number of auxiliary components^17,18^. A comprehensive analysis of existing literature reveals that the composition of reaction buffers differs widely between protocols, both in the number of optional components and in the concentrations of individual components (**Table S1**). These complex systems, while effective, often lead to high costs, laborious preparation procedures, and potential interactions among components, posing significant challenges for further optimization and standardization, thereby limiting their widespread adoption in resource-constrained laboratories.

In recent years, the field of eukaryotic CFPS has seen remarkable progress in system optimization and simplification^19,20^. Highly optimized human *in vitro* translation systems have demonstrated that high-efficiency protein synthesis can be achieved with a minimal number of core components^19^. Recognizing the potential of such systematic optimization approaches, we hypothesized that similar strategies could be equally applicable and beneficial for *E. coli* CFPS systems by thoroughly analyzing the functions of existing components and integrating insights from previous reports on prokaryotic protein synthesis mechanisms. This study aimed to develop a highly simplified yet efficient *E. coli* cell-free protein synthesis system (*e*CFPS). We conducted a systematic component reduction screening of traditional *e*CFPS systems, successfully streamlining the core components from thirty-five to just seven. The subsequent meticulous optimization of these seven key components not only maintained but even improved protein expression levels. Through these dual efforts—simplifying the reaction mixture and developing a high-quality, dialysis-free “fast lysate” preparation—we established a highly accessible and robust *e*CFPS platform which will accelerate diverse applications in life science research.

## Results

### Streamlining *e*CFPS: removal of dispensable components

To develop a more streamlined and efficient *e*CFPS system, we performed a systematic screening of auxiliary components commonly found in traditional prokaryotic CFPS protocols^17,21^. Our objective was to identify dispensable components while maintaining or enhancing protein synthesis efficiency, thereby simplifying system preparation and reducing costs. Starting with a comprehensive reaction mixture containing up to thirty-five components^17^, we iteratively evaluated the contribution of individual constituents through luciferase reporter assays. Throughout this optimization, essential core components such as creatine phosphate (CrP), creatine kinase (CrK), ATP, GTP, magnesium, and potassium were maintained in the base reaction mixture^3,8,19^. Our systematic approach allowed us to precisely determine the impact of each component on overall protein synthesis yield, leading to the identification of both dispensable and critical factors.

Through this systematic screening, we first identified several components that could be entirely removed from the reaction mixture without compromising protein synthesis efficiency (**Figure 1**). Dithiothreitol (DTT), a reducing agent commonly included in both transcription and translation systems to maintain protein sulfhydryl groups in a reduced state and prevent aggregation, was found to be unnecessary within our specific system, as its removal did not affect the final protein expression levels (**Figure 1A**). Similarly, cyclic adenosine 3′,5′-monophosphate (cAMP), a regulator implicated in transcriptional regulation^22–24^, was also found to be dispensable for efficient protein synthesis (**Figure 1B**). Furthermore, we systematically evaluated other auxiliary components, including the molecular crowder polyethylene glycol 8000 (PEG8000), used to enhance macromolecular crowding^25–29^; ammonium ions (NH_4_ ^+^), typically included as an osmolyte and for its role in maintaining protein stability and solubility in *e*CFPS reactions^29–33^; and folinic acid, which serves as a crucial cofactor for nucleotide synthesis^8,33^. Our results consistently showed that removing these components had a negligible impact on overall protein synthesis performance (**Figure 1C-E**). These findings collectively demonstrate that a substantial portion of the auxiliary components in traditional *e*CFPS protocols can be eliminated, paving the way for a more streamlined and cost-effective system.

**Figure 1.**
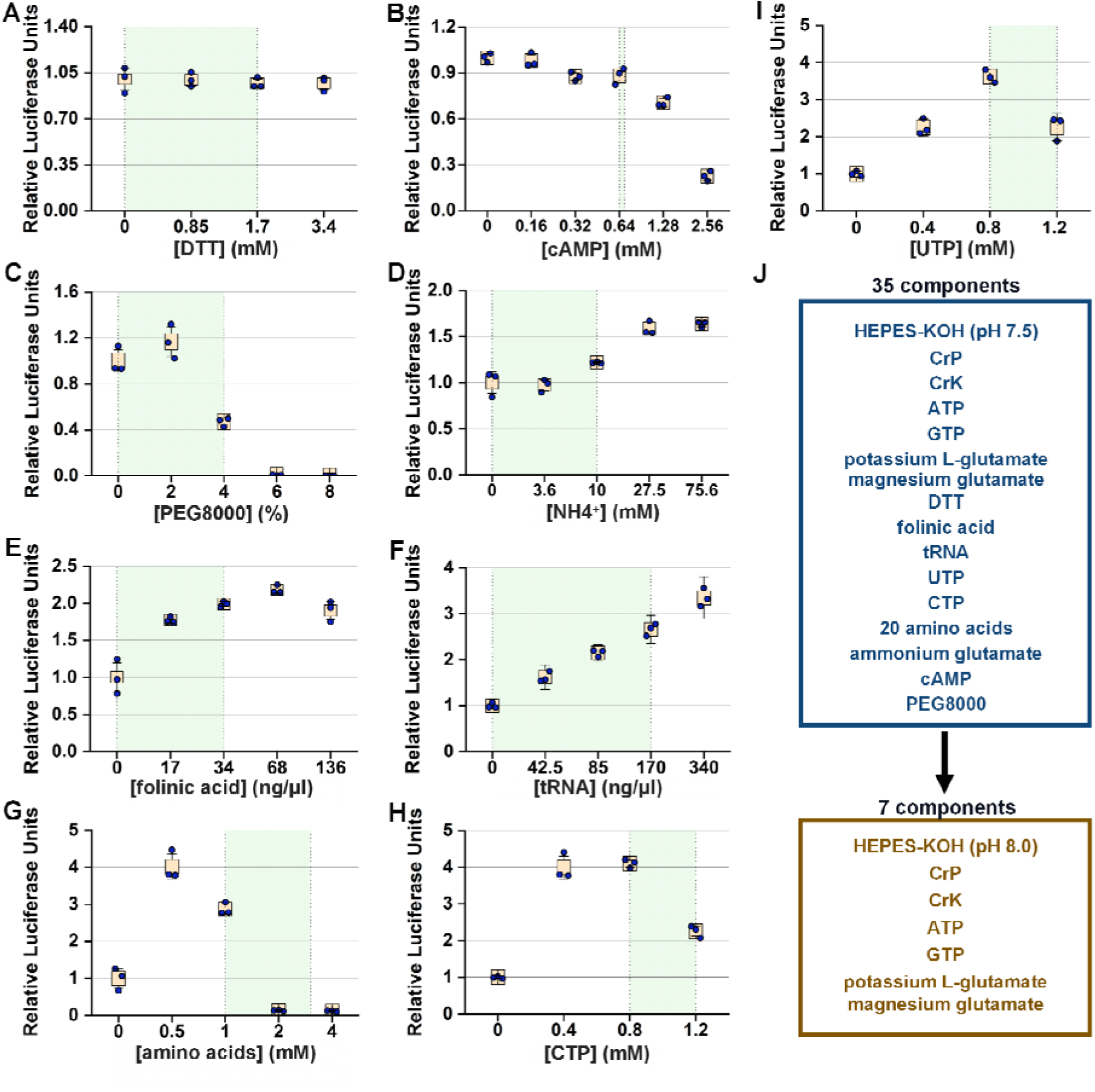
Optimization of *e*CFPS components. Protein expression levels from the *e*CFPS system were measured using an Nanoluciferase (NLuc) reporter DNA. Green area in the graphs indicate the common concentration range used in published protocols for *e*CFPS. Error bars represent the standard error (SE) of at least three independent reactions. (**A-E**) Protein expression levels of the *e*CFPS system supplemented with different concentrations of DTT (**A**), cAMP (**B**), PEG8000 (**C**), NH_4_^+^ (**D**), and folinic acid (**E**). (**F-I**) Protein expression levels of *e*CFPS with various concentrations of tRNA (**F**), amino acids (**G**), CTP (**H**) and UTP (**I**). (**J**) A summary of the supplement components before and after optimization.

An evaluation of various concentrations of amino acids and tRNA was conducted, as these components are fundamental building blocks for protein synthesis^34–36^. Previous studies on *e*CFPS have highlighted the rapid degradation of certain amino acids, such as arginine, cysteine, and tryptophan, necessitating their replenishment for prolonged synthesis^37,38^. Our results showed that while these components are critical for achieving high yields, protein synthesis still occurs in their absence (**Figure 1F-G**). This suggests that while they are optimizable, tRNA and amino acids are not strictly essential for the reaction to proceed, likely due to residual amounts within the cell lysate (**Figure 1F-G**). Furthermore, we evaluated the role of nucleoside triphosphates (NTPs) in our system, which are essential for coupled transcription-translation. Interestingly, we found that protein synthesis could proceed without the addition of CTP or UTP (**Figure 1H-I and Figure S1**). While adding either ATP or GTP alone resulted in a very weak reaction, the presence of both ATP and GTP together recovered the reaction to approximately 40% of the complete NTPs mix (**Figure S1B-C**). This highlights the critical role for ATPase-dependent chaperones (e.g., DnaK) and GTPase-dependent elongation factors (e.g., EF-Tu and EF-G)^39^, which are crucial for proper protein folding during synthesis^40^.

Ultimately, this comprehensive screening allowed us to successfully reduce the core reaction components from thirty-five to just seven (**Figure 1J**).

### Optimization of essential *e*CFPS components

While a core set of seven components was found to be sufficient for protein synthesis, we conducted further fine-tuning to maximize the system’s performance. First, we optimized the concentrations of key salts and energy components (**Figure 2**). Our findings reveal that the final protein expression level is highly dependent on the concentrations of both magnesium (Mg^2+^) and potassium ions (K^+^), which are fundamental for the structural integrity and catalytic activity of ribosomes, and various enzymatic reactions critical to *e*CFPS^41,42^. Through a detailed matrix-based optimization, we first identified the optimal concentrations of Mg^2+^ and K^+^ to achieve maximum protein expression (**Figure 2A**). Similarly, we conducted a separate screen to optimize the concentrations of Mg^2+^ and PEG8000 as previous reports have suggested a cooperative relationship between them^17^ (**Figure 2B**).

**Figure 2.**
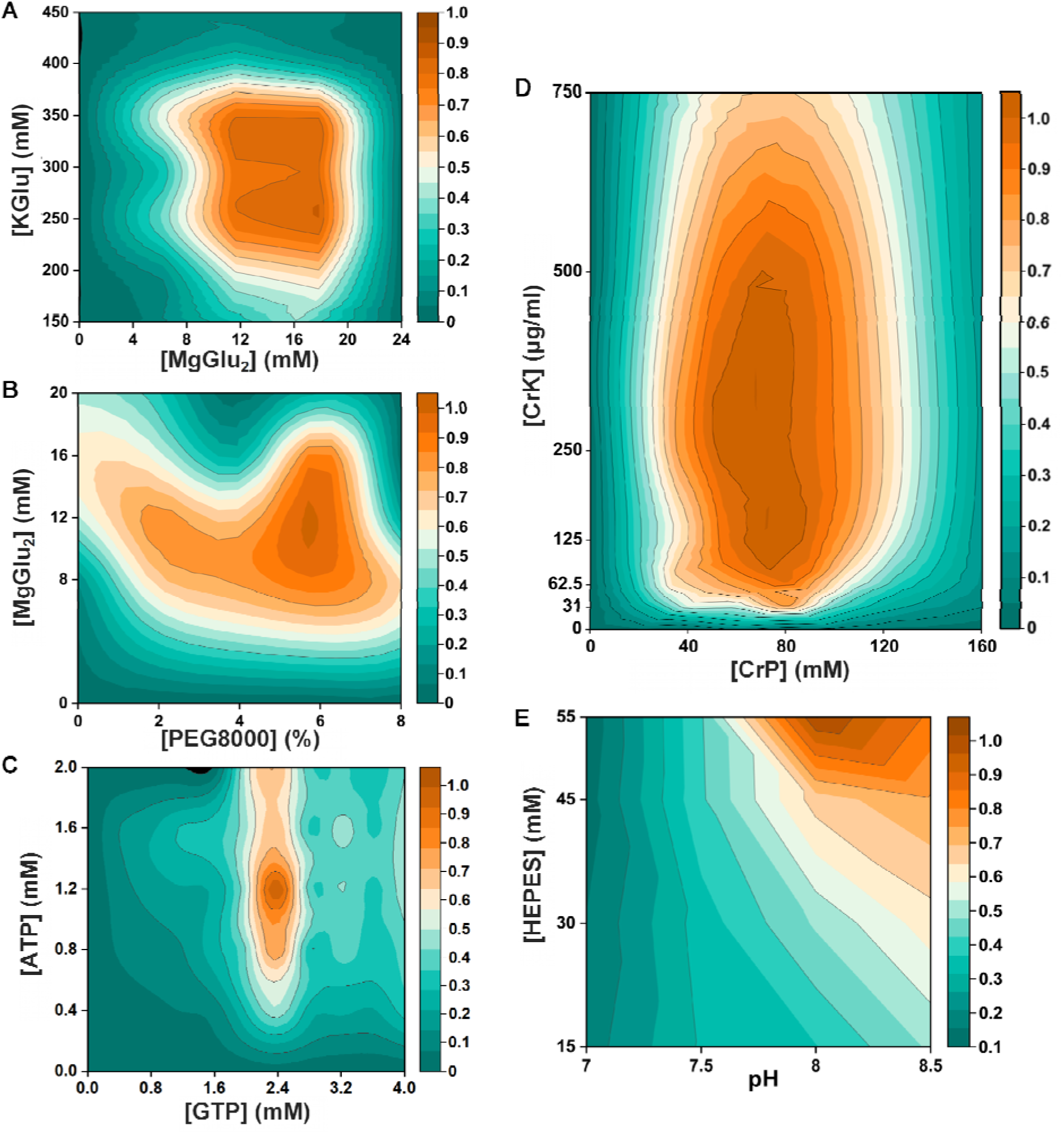
Optimization of essential components for *e*CFPS system. (**A**) Protein expression levels of the *e*CFPS system measured at varying concentrations of KGlu and MgGlu_2_. (**B**) Protein expression levels of the *e*CFPS system measured at varying concentrations of MgGlu_2_ and PEG8000. (**C**) Protein expression levels of the *e*CFPS system measured at varying concentrations of ATP and GTP. (**D**) Protein expression levels of the *e*CFPS system measured at varying concentrations of CrK and CrP. (**E**) Protein expression levels of the *e*CFPS system measured at varying pH and buffer concentrations. Data from all panels present mean ± SE, n = 3.

A comprehensive evaluation of the core energy components was also performed. We specifically focused on co-optimizing the concentrations of ATP and GTP, which serve as both energy sources and building blocks^43–45^, to maximize the yield of protein synthesis (**Figure 2C**). Additionally, the energy regeneration system, primarily composed of CrP and CrK, is vital for maintaining sustained ATP levels^46,47^. We found that CrP is crucial, with no reaction occurring in its absence (**Figure S2A**). Initial experiments showed that while CrK addition significantly boosted translation efficiency early on, reactions without exogenous CrK could achieve a protein expression level approximately four times higher at later time points (2 hours or more), suggesting the presence of an endogenous CrK-like enzyme in the lysate (**Figure S2B**). A co-optimization screen for CrP and CrK concentrations was also performed, and our results showed that optimizing the concentrations of these components is critical for achieving and sustaining high protein expression (**Figure 2D**). Furthermore, we optimized the pH and concentration of the HEPES buffer, finding that a specific range was critical for maintaining stable and high protein expression (**Figure 2E**). These multi-faceted optimizations ensured that each component was present at its ideal concentration, leading to a synergistic effect that significantly boosted the system’s overall performance.

### Performance characterization of the optimized *e*CFPS

To thoroughly evaluate the performance of our optimized *e*CFPS, we conducted a series of experiments. We first investigated the kinetics of protein synthesis measuring the expression over time at different temperatures (25°C, 30°C, and 37°C). Our results demonstrate that the system exhibits robust protein synthesis across this range, with the highest expression rates observed at the physiological temperature of 37°C (**Figure 3A**). Next, we assessed the system’s sensitivity to DNA concentration, finding that it could efficiently utilize a wide range of reporter DNA templates to produce quantifiable protein products, which were validated by both luminescence assays and western blot analysis (**Figure 3B**). Furthermore, we investigated the impact of varying cell lysate volume ratios, comparing our initial system with the newly optimized *e*CFPS. Our findings indicate that the optimized system maintains superior protein expression even at different lysate-to-buffer ratios (**Figure 3C**).

**Figure 3.**
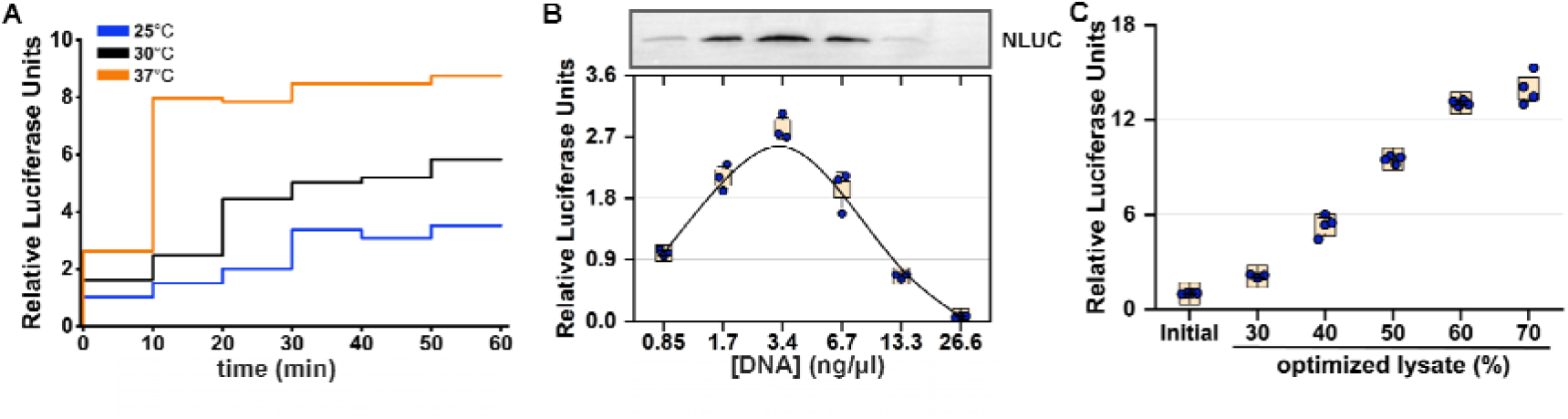
Characterization of the optimized *e*CFPS system. (**A**) Kinetics of protein synthesis at 25°C, 30°C and 37°C over a 60-minute period. Data present mean ± SE, n = 3. (**B**) Protein expression levels of the *e*CFPS system measured at varying DNA concentrations for a reporter encoding a FLAG-tagged NLuc. The protein product was quantified via a luminescence assay and confirmed by western blotting. Data present mean ± SE, n = 3. (**C**) Comparison of protein yield. The ‘initial’ system denotes the traditional 35-component reaction mixture prior to optimization, serving as a baseline control for benchmarking the streamlined system. Data present mean ± SE, n = 4.

To distinguish whether the increased protein expression in the optimized system was due to enhanced transcription or translation, we conducted assays using both DNA and pre-transcribed mRNA templates. The results (**Figure S3A**) indicate that while the translation efficiency in the streamlined system showed a slight decrease compared to the initial system, the significant improvement in transcription efficiency resulted in a higher overall protein yield. Direct quantification via RT-qPCR revealed a remarkable increase in transcript levels in our optimized system compared to the standard initial system (**Figure S3B**). Notably, even when the initial system was supplemented with excess T7 RNA polymerase (400 ng/µL), the accumulated transcript levels remained substantially lower than those of our optimized system. Furthermore, T7 RNA polymerase titration confirmed that the initial system saturates at 800 ng/µL —a concentration nearly ten times higher than standard protocols (∼90–100 ng/µL) — while remaining ∼45-fold less productive than our optimized system (**Figure S3C**).

### Benchmarking and expression of challenging proteins

We benchmarked our system against two widely used systems: the classical Phosphoenolpyruvate (PEP)-based system and our 35-component initial CrP/CrK-based system^6,7^ (**Figure 4**). We utilized NLuc, representing a broadly applicable protein, and super-folder green fluorescent protein (sfGFP), with codons optimized for *E. coli* as reporter proteins and monitored their expression over time (**Figure 4A-B, Figure S4A**). Our results show that the optimized *e*CFPS consistently outperforms both the classical and initial systems, achieving significantly higher protein expression levels in a shorter period (**Figure 4A-B**). The robust expression was further confirmed by western blot analysis, which provided clear visual evidence of the superior protein yield from our optimized system (**Figure 4C-D, Figure S4B-C**). Finally, we validated the function of our system through antibiotic-mediated inhibition assays. Our results demonstrate that protein expression in the optimized *e*CFPS can be effectively inhibited by standard antibiotics, confirming its robust and native-like translation machinery (**Figure S4D**).

**Figure 4.**
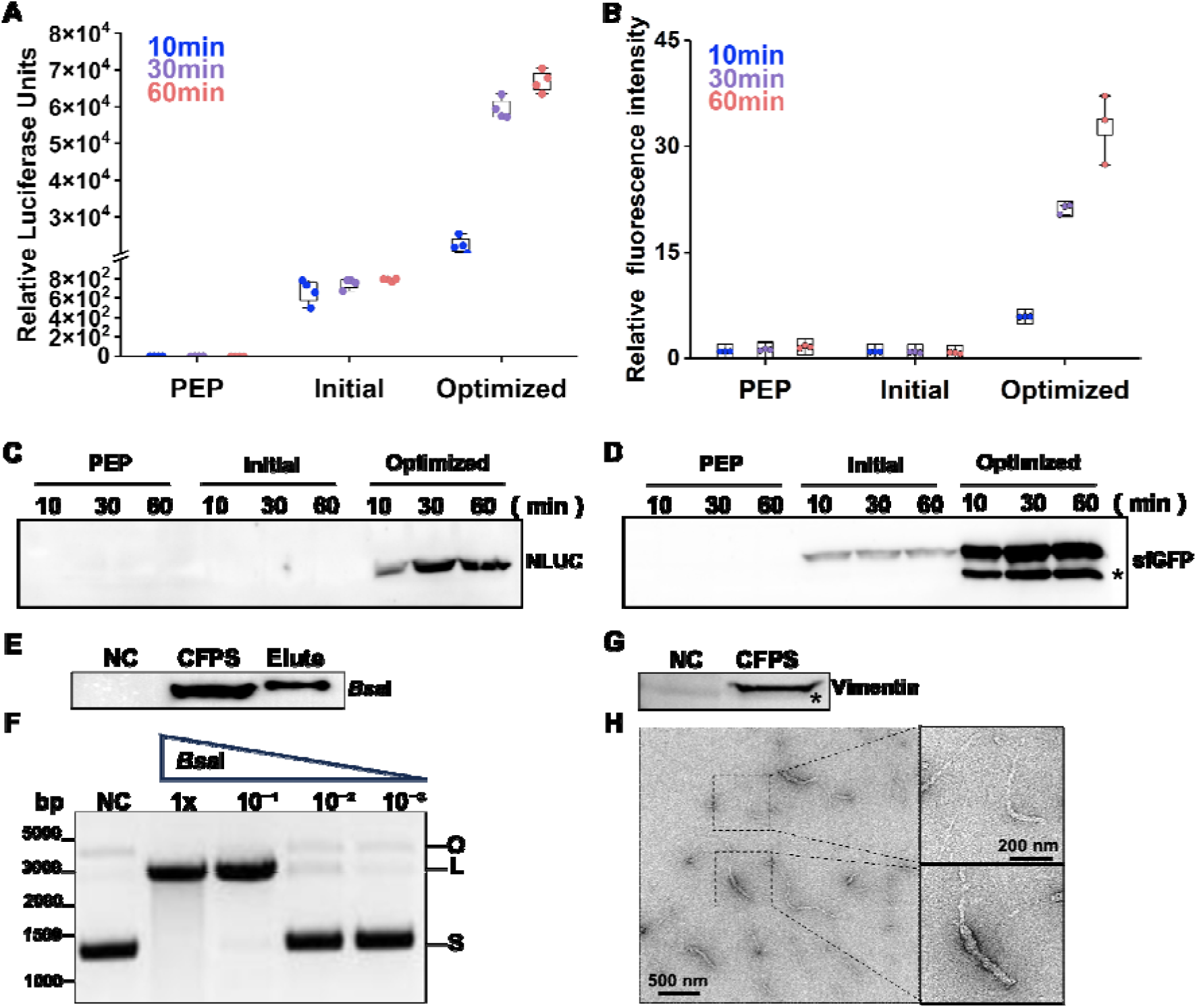
Benchmarking the optimized *e*CFPS system with different DNA templates. **(A)** NLuc protein expression kinetics over time, comparing the PEP-based, initial CrP/CrK-based, and optimized CrP/CrK-based energy regeneration systems. Data present mean ± SE, n = 4. (**B**) sfGFP protein expression kinetics over time from the three energy regeneration systems. Data present mean ± SE, n = 3. (**C-D**) Western blot validation of protein expression for NLuc (**C**) and sfGFP (**D**) from the different *e*CFPS system shown in (**A-B**). Protein products were detected using an anti-FLAG antibody. The asterisk (*) indicates a non-specific band. (**E**) Western blot detection of His-FLAG-*Bsa*I expressed by the optimized *e*CFPS system using an anti-FLAG antibody. (**F**) Agarose gel electrophoresis confirming the functional activity of *e*CFPS-synthesized *Bsa*I via cleavage of a substrate plasmid. A 10-fold serial dilution of *Bsa*I was with 1x representing 0.05 mg/mL. NC (negative control) indicates no plasmid in the *e*CFPS reaction. S, L, and O indicate the respective position of the supercoiled, linear, and open circular forms of the plasmid. (**G**) Western blot analysis of vimentin expressed by the optimized *e*CFPS system using an anti-vimentin antibody. (**H**) Negative-stain electron microscopy image showing that vimentin expressed via *e*CFPS can successfully self-assemble into filaments *in vitro*.

To further validate the system’s capability in synthesizing challenging, functional proteins, we focused on the restriction endonuclease *Bsa*I (**Figure 4E-F**). Restriction endonucleases like *Bsa*I are notoriously difficult to express *in vivo* due to their specific DNA-cleaving activity, which is highly cytotoxic to the host *E. coli* cells. Traditional recombinant expression requires the compulsory co-expression of a corresponding methylase to protect the host genome, adding significant complexity to the system^48,49^. Leveraging the inherent cell-free advantage of circumventing cytotoxicity, we first demonstrated the system’s utility by synthesizing the restriction endonuclease *Bsa*I^50^, confirming its specific enzymatic activity via a DNA cleavage assay, which underscores the system’s ability to produce complex, functional enzymes (**Figure 4E-F**). We confirmed that the *e*CFPS synthesized *Bsa*I was functionally active, successfully cleaving its substrate plasmid DNA (**Figure 4F**). Based on the standard definition of enzyme activity (1 U digests 1 μg of substrate DNA in 20 μL at 37°C in 1 h), we calculated the enzyme activity of our *e*CFPS-produced *Bsa*I to be in the range of 2×10^3^ ∼ 2×10^4^ U/mg (**Figure 4F**). The efficient production of this active enzyme demonstrates the power of our simplified system for producing complex, cytotoxic proteins.

Next, we successfully expressed vimentin, an intermediate filament protein (**Figure 4G**). Vimentin is a type III intermediate filament protein that forms part of the cytoskeleton^51^. It is known to be difficult to express and handle due to its high propensity for aggregation. Our *e*CFPS system efficiently expressed vimentin **(Figure 4G).** Crucially, the vimentin expressed in our CFPS system demonstrated successful *in vitro* self-assembly into filaments (**Figure 4H**), which was confirmed by negative-stain electron microscopy, confirming the system’s robust performance even for difficult-to-express, aggregation-prone proteins (**Figure 4E-H**). We also benchmarked the absolute productivity of our optimized system against a high-end commercial cell-free system using sfGFP (**Figure S4G**). The optimized system achieved an absolute yield of 0.46 mg/mL, significantly outperforming the commercial alternative (approximately 0.21 mg/mL).

### A simplified method for bacterial lysate preparation and quality control

In addition to optimizing the reaction buffer, we sought to simplify the entire *e*CFPS procedure by developing an easy-to-use method for preparing the bacterial cell lysate. During lysate making, traditional methods, which often rely on time-consuming runoff and dialysis procedures^21,8^, were replaced with a high-pressure homogenizer for efficient cell disruption, eliminating the need for additional dialysis. Compared to traditional biochemical methods such as ultrasonication, lysozyme treatment, or freeze-thaw cycles^52^, this approach is faster and more convenient (**Figure 5A**). Additionally, endogenous T7 RNA polymerase in the optimized *e*CFPS lysate obviates exogenous addition, simplifying preparation and reducing costs while maintaining high translation efficiency (**Figure 5A**). We validated this new “fast lysate” preparation by rigorous quality control. We tested lysates prepared from cells harvested at various optical densities at 600 nm (OD_600_). While the density did not significantly impact protein expression levels, a harvest optical density of OD_600_=2 yielded the best performance (**Figure 5B**). The quality of the lysate, particularly the integrity of its translational machinery, was further confirmed by sucrose gradient centrifugation (**Figure S5A**) and negative-staining transmission electron microscopy (TEM) images (**Figure S5B**) showing well-resolved 70S ribosomes.

**Figure 5.**
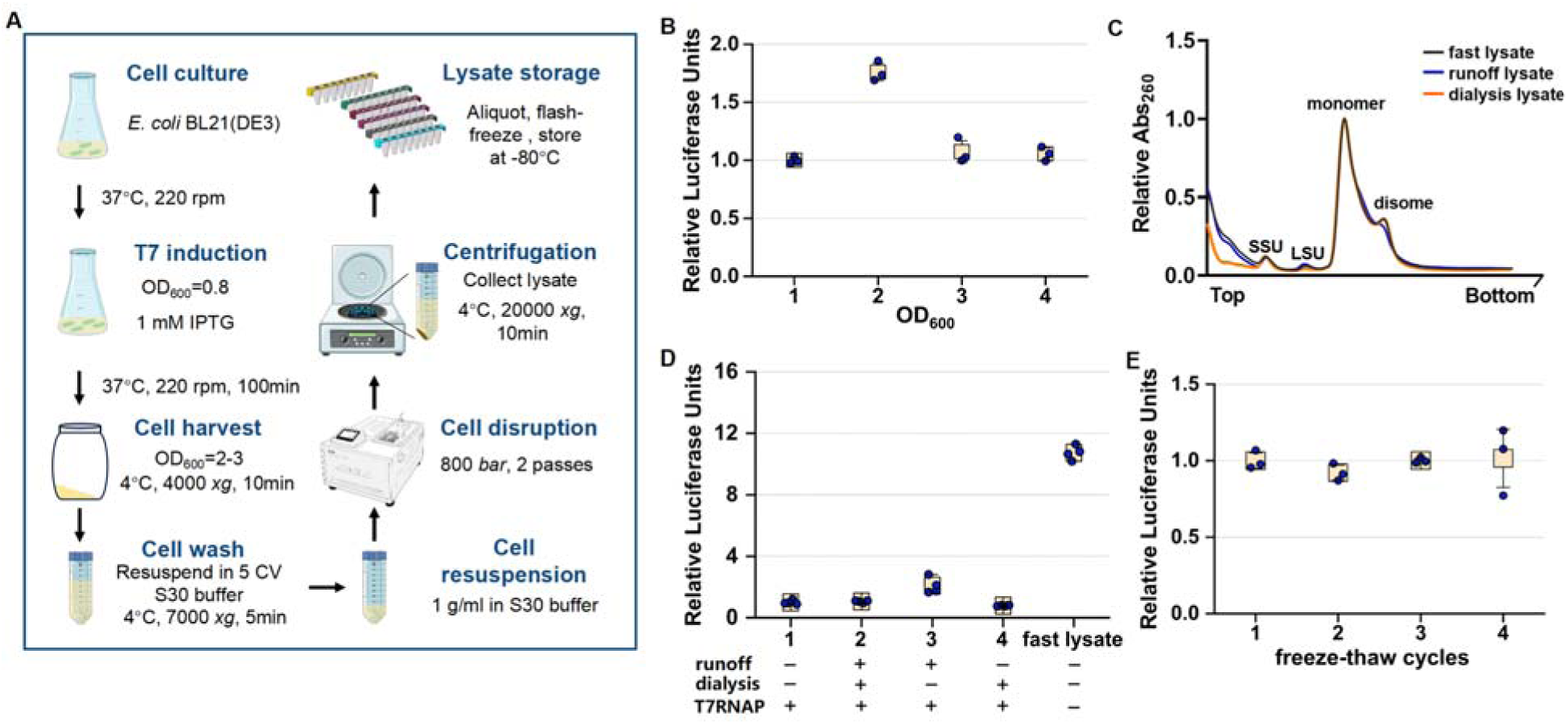
Preparation of *e*CFPS from cultured *E. coli* Cells. (**A**) Flowchart of the *e*CFPS preparation procedures. (**B**) Comparison of reaction efficiency in *e*CFPS using lysate with bacteria cells harvested at different optical density. Data present mean ± SE, n = 3. (**C**) Sucrose gradient sedimentation analysis of different lysates used for *e*CFPS, revealing the presence of ribosome monomers. (**D**) Comparison of protein expression levels in *e*CFPS system using lysates prepared by runoff, dialysis, and rapid endogenous T7 RNA polymerase induction. Data present mean ± SE, n = 4. (**E**) Comparison of reaction efficiency in *e*CFPS using lysates after different numbers of freeze-thaw cycles. Data present mean ± SE, n = 3.

To further simplify the protocol, we evaluated the necessity of the traditional runoff and dialysis steps^8^ (**Figure 5A**). Our results from sucrose gradient analysis and TEM imaging confirmed that ribosomes remained intact, and the *e*CFPS expression signal was significantly higher without them (**Figure 5C-D, Figure S5B**). These findings confirm that both the runoff and dialysis steps are dispensable for our system, allowing for a significantly simplified and faster preparation. Additionally, our reliance on endogenously expressed T7 RNA polymerase yields a significantly higher expression signal, outperforming previously reported *e*CFPS systems by up to four-fold (**Figure 5D**).

Our optimized lysate preparation method yielded highly active and stable extracts. It is capable of sustaining protein synthesis for up to 120 minutes (**Figure S5C**) and maintains protein synthesis efficiency even after multiple freeze-thaw cycles, making it highly suitable for routine laboratory use (**Figure 5E**).

## Discussion

This study presents a streamlined and highly efficient *e*CFPS platform achieved by reducing the reaction components from 35 to 7. The performance of this simplified system was evaluated against the initial 35-component setup and PEP-based platforms, which serve as established reference baselines. The improvement in protein yield over these comparative standards, coupled with stable absolute productivity, supports the efficacy of the optimized reaction buffer and lysate preparation.

A notable finding was the ability of our system to function effectively without several components traditionally considered essential, which we attribute to the metabolic capacity of the fast lysate. By bypassing traditional runoff and dialysis steps, this protocol preserves active endogenous enzymes capable of synthesizing building blocks from residual precursors. For instance, functional pathways for synthesizing specific amino acids, such as deriving Cys and Trp from Ser, and generating Asn and Gln from Asp and Glu, are maintained^53^. Similarly, endogenous nucleotide metabolic enzymes, powered by the CrP/CrK regeneration system, convert residual pools into functional CTP and UTP to support coupled transcription and translation^54^. Further analysis indicates that the performance gain is primarily driven by enhanced transcription, which more than compensates for a modest reduction in translational efficiency in the streamlined environment. This improvement does not stem from a simple increase in T7 RNA polymerase availability; instead, it reflects a systemic synergy where the streamlined buffer and the optimized metabolic environment of the lysate together alleviate transcriptional bottlenecks inherent in traditional platforms. This systemic rescue of transcription is directly supported by our RT-qPCR analysis, which confirmed that relieving buffer complexity, rather than merely increasing polymerase dosage, is key to overcoming transcriptional bottlenecks. Ultimately, this metabolic self-sufficiency, combined with the preservation of critical endogenous factors through the omission of dialysis, maintains the lysate in a highly active state that permits significant simplification of the reaction chemistry while driving superior performance.

The applicability of the optimized *e*CFPS was tested through the synthesis of targets beyond standard reporters. The production of active *Bsa*I restriction enzyme, a cytotoxic and difficult-to-express protein, and the functional assembly of vimentin, a difficult-to-handle intermediate filament protein^51^, further validate the superior robustness and translational quality of our optimized system.

Despite its efficiency, certain limitations are acknowledged. As a prokaryotic platform, it lacks the eukaryotic chaperone systems (e.g., Hsp70/Hsp90 families) and post-translational modification (PTM) machinery required for many human proteins^10^. While we demonstrated that DTT is dispensable for the targets tested here, specialized cysteine-rich proteins may still require exogenous reducing agents to ensure proper folding. Furthermore, while the fast lysate protocol significantly lowers the barrier to entry, the expression of certain membrane proteins will likely still require the systematic screening of detergents or synthetic lipids to ensure proper integration and stability. In conclusion, our optimized *e*CFPS system reduces reaction complexity while improving performance. By simplifying both the buffer composition and the lysate preparation procedure, we have established a cost-effective, and user-friendly platform. This system is expected to facilitate high-throughput protein engineering, compound screening, and diagnostic development across the broader scientific community.

## Methods

### Plasmid construction

All plasmids and primers used in this study are detailed in **Table S2**. The NLuc and sfGFP reporter genes, each fused with a FLAG-tag, were cloned into a T7-driven expression vector. The sfGFP gene was optimized for *E. coli* codon usage to ensure efficient translation. The vector backbone was constructed using standard molecular biology techniques.

The *Bsa*I linear DNA templates was constructed from three DNA fragments amplified by PCR. To enhance template stability and mitigate nuclease degradation within the CFPS system, approximately 300 bp sequences were incorporated upstream and downstream of the *Bsa*I coding sequence. The fragments encoding the T7 promoter/upstream sequence (Fragment 1) and the T7 terminator/downstream sequence (Fragment 2) were amplified from the pJL1 plasmid. Separately, the gene fragment for the restriction endonuclease *Bsa*I was amplified from the pUC_*Bsa*I plasmid (Fragment 3). These three fragments were subsequently fused using overlap PCR to generate the final linear template, which was confirmed to lack the recognition sequence for the target *Bsa*I enzyme.

### *E. coli* cell culture and lysate preparation

*E. coli* S30 cell lysate for CFPS was prepared using a simplified protocol adapted from a previously reported method^8^. Briefly, *E. coli* BL21(DE3) was cultured in LB medium at 37°C with shaking. Once the OD_600_ reached 0.8, endogenous T7 RNA polymerase expression was induced with 1 mM IPTG. The culture was then grown for an additional 2 hours before being harvested (when the OD_600_ reached 2-3). The cells were collected by centrifugation at 4000 *g* for 10 minutes at 4°C. The cell pellet was washed with 5 volumes of cold S30 buffer (10 mM HEPES-KOH pH 7.5, 14 mM Mg(OAc)_2_, 60 mM KOAc) and then resuspended in S30 buffer at a ratio of 1 mL per gram of wet cell paste. Cell lysis was performed using a high-pressure homogenizer at 800 *bar* for two passes. The lysate was then clarified by centrifugation at 20,000 *g* for 10 minutes at 4°C, and the resulting supernatant, designated as the “fast lysate”, was collected without subsequent runoff or dialysis steps. This omission ensures maximal retention of endogenous components, contributing directly to the observed system efficiency. Lysate quality control was assessed by (i) sucrose density gradient analysis and negative-staining TEM to confirm the integrity of 70S ribosomes, and (ii) functional testing by monitoring the synthesis and activity of the challenging restriction enzyme, *Bsa*I and vimentin. Finally, the fast lysate was aliquoted and flash-frozen in liquid nitrogen before being stored at -80°C until use.

A step-by-step protocol is provided in the supplementary protocol.

### *e*CFPS reactions for luciferase reporter assay

The optimized *e*CFPS reaction was performed in a 10 μL total volume containing the components listed in **Table S3** and **Table S4**. Plasmid of NLuc reporter gene was used as the template.

### Quantitative Real-Time PCR (RT-qPCR)

To evaluate the transcriptional levels of the NLuc reporter gene across different cell-free systems, quantitative real-time PCR (RT-qPCR) analysis was performed.

Each reaction was carried out in a total volume of 200 μL with a template plasmid concentration of 3.4 ng/μL. The reactions were incubated at 37 °C for 30 min. A parallel 0 min reaction control was established as the baseline calibrator. Total RNA was extracted using the TRIzol reagent (Invitrogen) according to the manufacturer’s protocol, and the extracted RNA pellet was dissolved in an equal volume (900 μL) of nuclease-free water.

To synthesize cDNA, an equal volume (2 μL) of total RNA from each sample was reverse transcribed using the HiScript III 1st Strand cDNA Synthesis Kit (Vazyme, R312). To thoroughly rule out plasmid or genomic DNA contamination, no-template controls (NTC) and no-reverse transcriptase (no-RT) controls for all systems were prepared and analyzed in parallel. Quantitative real-time PCR was performed using the ChamQ SYBR Color qPCR Master Mix (Vazyme, Q411) on an Applied Biosystems Real-Time PCR System (Thermo Fisher Scientific) according to the manufacturer’s instructions. The relative fold changes of the NLuc transcript levels were calculated using the comparative 2^-ΔΔCt^ method, with the 0 min sample of each group serving as the calibrator. All assays were performed with three independent biological replicates.

### CFPS expression, protein purification and analysis

Protein expression levels were quantified using NLuc and sfGFP reporter systems. NLuc activity was measured with the Nano-Glo luciferase assay system (Promega). Reaction mixtures were diluted as specified in the source data to prevent signal saturation and then analyzed on a microplate luminometer (BERTHOLD, Centro XS^3^ LB 960) to obtain the luminescence units as described previously^55,56^. Relative Luciferase Units (RLU) were determined by normalizing the raw luminescence intensity of each experimental group to that of a designated control group (typically the ‘initial’ 35-component system). This normalization was performed by dividing the absolute signal of the sample by the mean signal of the control, allowing for consistency across different experimental batches and measurement sessions. SfGFP fluorescence was used to quantify protein synthesis following the methods reported^57^. Briefly, 5 µL of the reaction mixture was added to 195 µL of fluorescence assay buffer (20mM HEPES-KOH pH7.5, 100mM NaCl, 5mM magnesium glutamate). Fluorescence was measured on a Cell Imaging Multimode Reader (BioTek, Cytation 5) with an excitation wavelength of 485 nm and an emission wavelength of 528 nm.

Western blotting analysis was performed to confirm protein synthesis of the *e*CFPS reactions. Synthesized proteins were detected using a primary anti-FLAG (Sigma, Cat#: M185-3L). Chemiluminescence was detected using a Gel Imaging System (Tanon).

### Preparation of a sfGFP standard curve

To establish a standard curve for the absolute quantification of sfGFP yield, *E. coli* BL21(DE3) transformed with the pJL1-sfGFP plasmid was inoculated into 5 mL of LB medium containing kanamycin and cultured overnight. Cells were harvested by centrifugation (10,000 *g*, 5 min), resuspended in lysis buffer (50 mM Tris-HCl pH 7.5, 300 mM NaCl), and disrupted using a high-pressure homogenizer at 800 *bar* for two passes. The clarified supernatant (10,000 *g*, 20 min) was purified via His-tag affinity chromatography, using 30 mM and 300 mM imidazole for washing and elution, respectively, with purity confirmed by SDS-PAGE.

The purified sfGFP was quantified via an Enhanced BCA Protein Assay Kit (Beyotime) and serially diluted in 50 mM HEPES buffer (pH 7.5) to concentrations ranging from 0.042 to 0.68 mg/mL. Fluorescence was measured using a Cytation 5 Multi-Mode Microplate Reader (BioTek) at an excitation of 485 nm and emission of 528 nm in flat-bottom 96-well half-area black plates. Each dilution was tested in triplicate to generate a standard curve for converting fluorescence intensity into absolute protein concentration (mg/mL).

### Expression and purification of vimentin and *Bsa*I

The gene for human vimentin (GenBank NP_003371.2) was sub-cloned into the pET507a vector for expression. *Bsa*I (4 mL reaction) and vimentin (1 mL reaction) proteins were synthesized using a CFPS system. The core reaction mixture for both proteins consisted of 16 mM magnesium glutamate, 250 mM potassium glutamate, 1.2 mM ATP, 2.4 mM GTP, 80 mM CrP, 125 μg/ml CrK, 13.3 ng/μl or 7.2 ng/μl DNA template, and 50% cell extract.

For *Bsa*I purification, the 4 mL reaction mixture was centrifuged at 20,000 *g*, and the supernatant was incubated with 600 μl of Ni beads for 1 h at 4°C. The beads were washed five times with 2 mL of wash buffer (40 mM HEPES-KOH pH 7.5, 50 mM imidazole, 500 mM NaCl). Elution was performed with 2 mL of elution buffer (40 mM HEPES-KOH pH 7.5, 500 mM imidazole, 500 mM NaCl). The eluted *Bsa*I was desalted using a column (desalting buffer: 40 mM HEPES-KOH pH 7.5, 500 mM NaCl, 1 mM DTT), concentrated to 1 mg/ml, and stored at -80°C in the presence of 10% glycerol. *Bsa*I enzymatic activity was validated in a 20 μL reaction containing 2 μL of 10x rCutsmart (NEB), 2 μg of a substrate plasmid, and *Bsa*I enzyme diluted in a 10-fold gradient. The reaction was incubated at 37°C for 1 h and visualized via agarose gel electrophoresis.

Vimentin synthesis was carried out in a 5 mL nuclease-free tube with static incubation at 37°C for 13 h. The *Bsa*I synthesis was performed in a 10 mL nuclease-free tube at 30°C with shaking at 150 rpm for 13 h.

### Vimentin filament assembly

Vimentin, expressed via CFPS, was refolded and prepared for assembly through a multi-step gradient dialysis protocol. The protein was dialyzed at room temperature in a buffer (5 mM Tris−HCl pH 8.5, 1 mM EDTA, 0.1 mM EGTA, 5 mM β-ME) sequentially containing 6 M, 4 M, and 2 M urea, with each step lasting 30 min. Subsequent overnight dialysis was performed at 4°C in the same buffer lacking urea. The protein was finally dialyzed for 1 h at room temperature into 5 mM Tris pH 8.5, 5 mM β-ME, ensuring the protein was in its tetramer configuration, after which the sample was concentrated. To initiate filament assembly, 20 μl of the concentrated protein solution was quickly mixed with 20 μl of assembly buffer (5 mM Tris pH 8.5, 170 mM NaCl, 100 mM KCl, 5 mM MgCl) in a 200 μl PCR tube. The assembly reaction was incubated for 1 h in a PCR thermocycler preheated to 25°C before being immediately transferred to ice. Filament formation was subsequently verified by negative-stain transmission electron microscopy.

### Negative-staining transmission electron microscopy (TEM)

Samples from vimentin assembly, the TEM samples were prepared by negative staining. A glow-discharged, carbon-coated copper grid was first rinsed with 3 μL of protein buffer (5 mM Tris pH 8.5, 5 mM β−ME). Then, 3 μL of the assembled sample solution was incubated on the grid for 20s. The sample was fixed using 3 μL of 0.8% glutaraldehyde solution for 20s. Finally, the grid was stained twice with 3 μl of uranyl acetate solution, with each staining step lasting 90 s. Samples were observed using a Talos F200C transmission electron microscope at a nominal magnification of 92,000.

Samples from the SDG were prepared for TEM. Fresh, glow-discharged carbon-coated grids were immersed in 3 μL of the corresponding sample for 2 minutes. Excess solvent was removed, and the grids were stained with uranyl acetate for 1 minute and air dried.

### Sucrose density gradient (SDG) analysis

A volume of 170–180 μL of lysate was loaded onto a 10%–40% (w/w) SDG. The gradient buffer contained 10 mM HEPES-KOH (pH 7.5), 14 mM Mg(OAc)_₂_, 60 mM KOAc. The gradient was prepared using a simple diffusion method (Biocomp) and centrifugation at 38,000 rpm for 2.5 hours at 4°C in a Beckman SW41Ti rotor. Following centrifugation, the gradient was fractionated using a density gradient fractionation system (Biocomp), with continuous monitoring of the absorbance at 260 nm.

## Acknowledgements

We thank Alexey Amunts for helpful discussions. We also thank Jian Li and Wanqiu Liu from Shanghai Tech University for providing the endonuclease plasmids and sharing the linear fragment construction method. This work was supported by grants from the National Key Research and Development Program of China (2022YFA0807100), National Natural Science Foundation of China (32171291 and 32371351), Shandong Basic Research Special Zone (Basic Medical Sciences YDZX2024154 and YDZX2025154), Nature Science Foundation of Shandong Province (ZR2021QC002) the Shandong Excellent Young Scientists Fund Program (2022HWYQ-025), Taishan Scholars Program (tsqnz20221104), Cutting Edge Development Fund of Advanced Medical Research Institute (GYY2023QY01) and the Cheeloo Youth Program of Shandong University to W.L..

## Contributions

X.L.,C.Z. and J.L. were responsible for obtaining experimental data and performing data processing. X.L. and C.Z. conducted biochemical experiments. X.L., C.Z., Z.Z. and W.L. analyzed the data and created charts. W.L. wrote the initial draft of the paper. All authors edited the paper.

## Notes

### Competing Interest Statement

The authors have declared no competing interest, with the exception of a pending patent application related to the optimized eCFPS system, which has been filed by the authors' institution, Shandong University.

### Summary of Updates

This revised version includes additional qPCR validation experiments and comparisons with commercial cell-free protein synthesis systems. The figures and supplementary information have also been updated accordingly.

