## Supplemental figures for "A simplified and highly efficient cell-free protein synthesis system for prokaryotes"

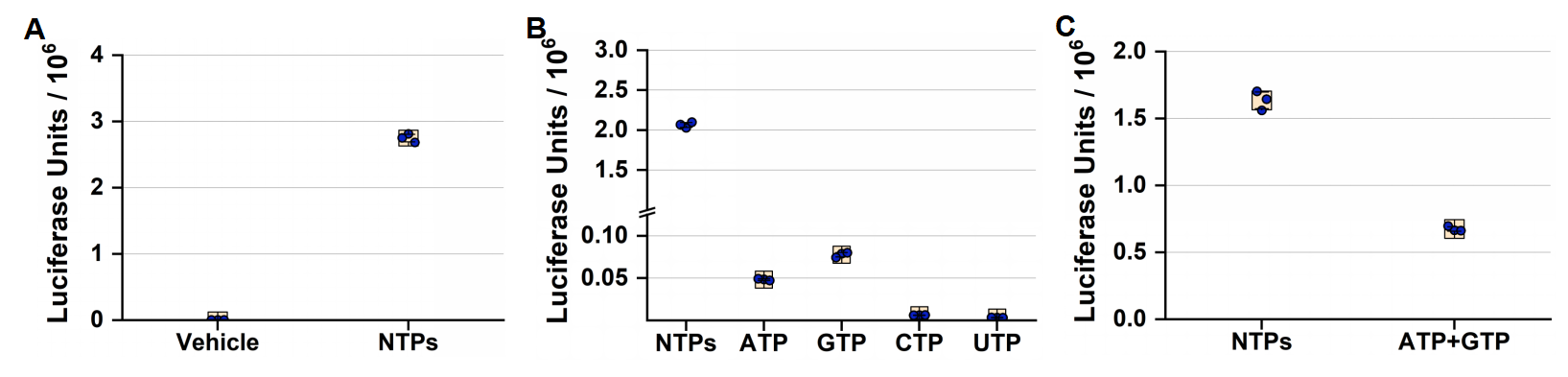

**Figure S1. The roles of NTPs in the reaction efficiency of *E. coli* cell-free protein synthesis (*e*CFPS). (A)** Reaction efficiency of *e*CFPS supplemented with and without NTPs. **(B)** Reaction efficiency of *e*CFPS supplemented with a complete mix of NTPs or individual NTPs (ATP, GTP, CTP or UTP). **(C)** Reaction efficiency of *e*CFPS supplemented with a mix of NTPs or a combination of ATP and GTP. Data present mean ± SE, n=3.

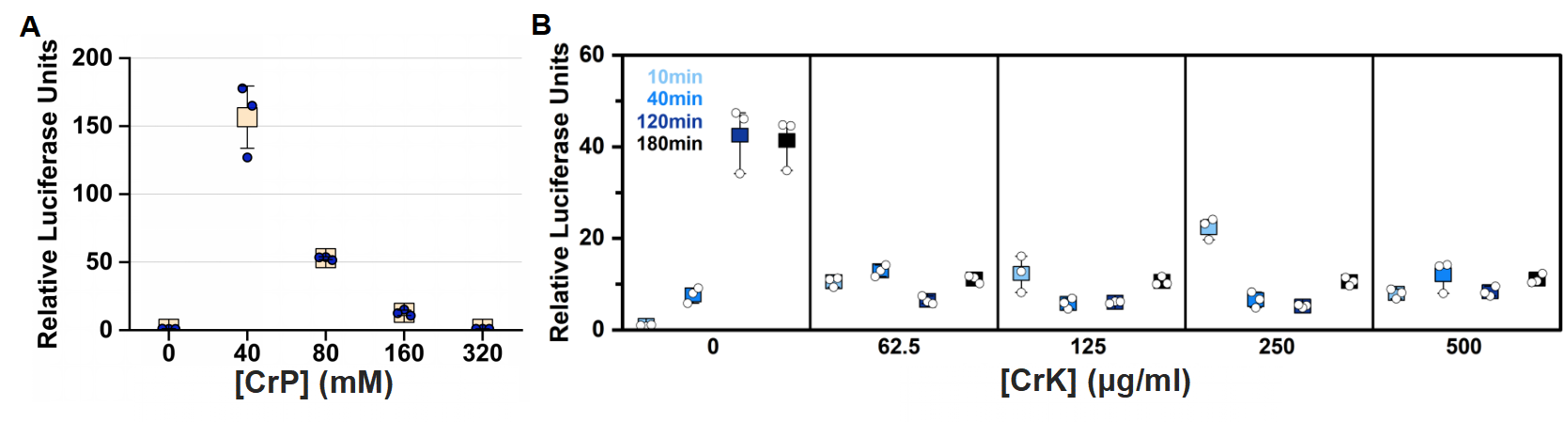

**Figure S2. Effects of CrK and CrP on the reaction efficiency of *e*CFPS. (A)** The reaction yield of optimized *e*CFPS at different concentrations of CrP. **(B)** Reaction efficiency of *e*CFPS at varying CrK concentrations and different reaction times. Data present mean ± SE, n=3.

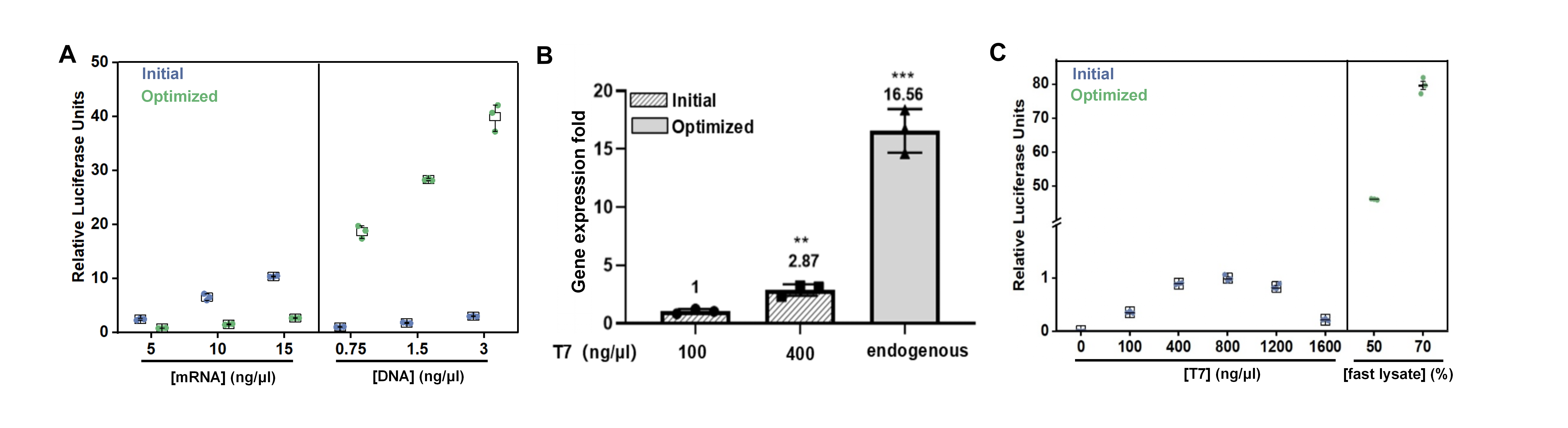

**Figure S3.** Performance evaluation and mechanistic analysis of transcriptional and translational efficiency. **(A)** Comparative evaluation of the initial and optimized systems using three different concentrations of DNA and mRNA templates to evaluate the transcription and translation efficiency. **(B)** Quantitative RT-qPCR analysis of reporter transcript levels in the initial and optimized systems. **(C)** Comparison between the initial system supplemented with varying concentrations of T7 RNA polymerase (0–1600 ng/μL) and the optimized system. Data present mean ± SE, n = 3.

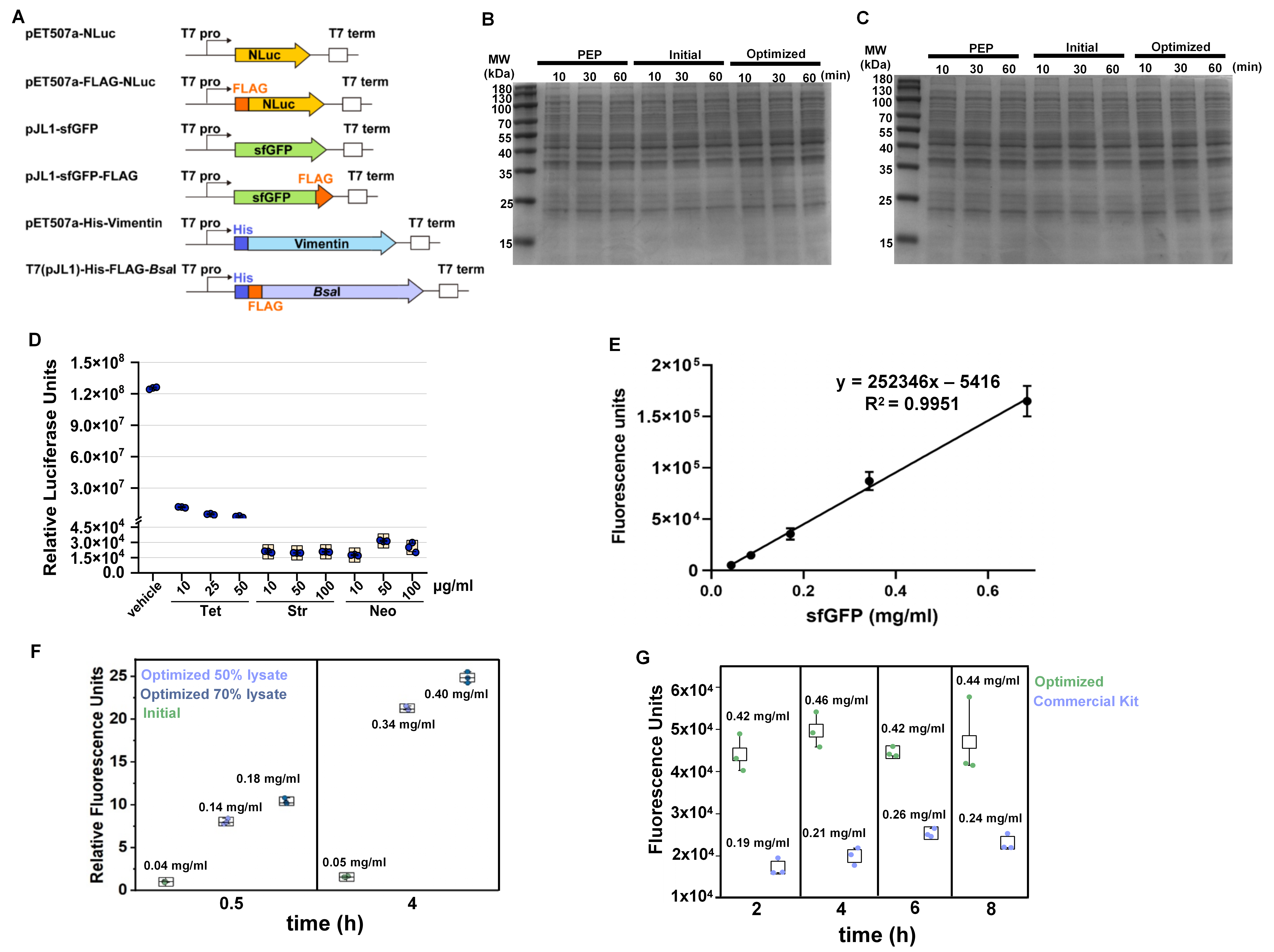

**Figure S4. Applications of *e*CFPS.** **(A)** Schematic overview of the vectors or expression cassettes used in this study, as described in the Methods section. **(B-C)** Total protein analysis by SDS-PAGE. SDS-PAGE gels for *e*CFPS reactions using NLuc **(B)** and sfGFP **(C)** as DNA templates. The SDS-PAGE results are shown as loading controls for western blot analysis of Figure 4. **(D)** Validation of antibiotic-mediated reaction inhibition using the *e*CFPS system. **(E)** Standard curve correlating sfGFP fluorescence intensity with absolute protein yield. **(F)** Quantitative comparison of **sfGFP protein yields** between the initial and optimized systems following short (0.5 h) and long (4 h) incubation. **(G)** Benchmarking of absolute protein productivity for the optimized system against a high-end commercial cell-free system at different time points using sfGFP as a reporter. Data present mean ± SE, n = 3.

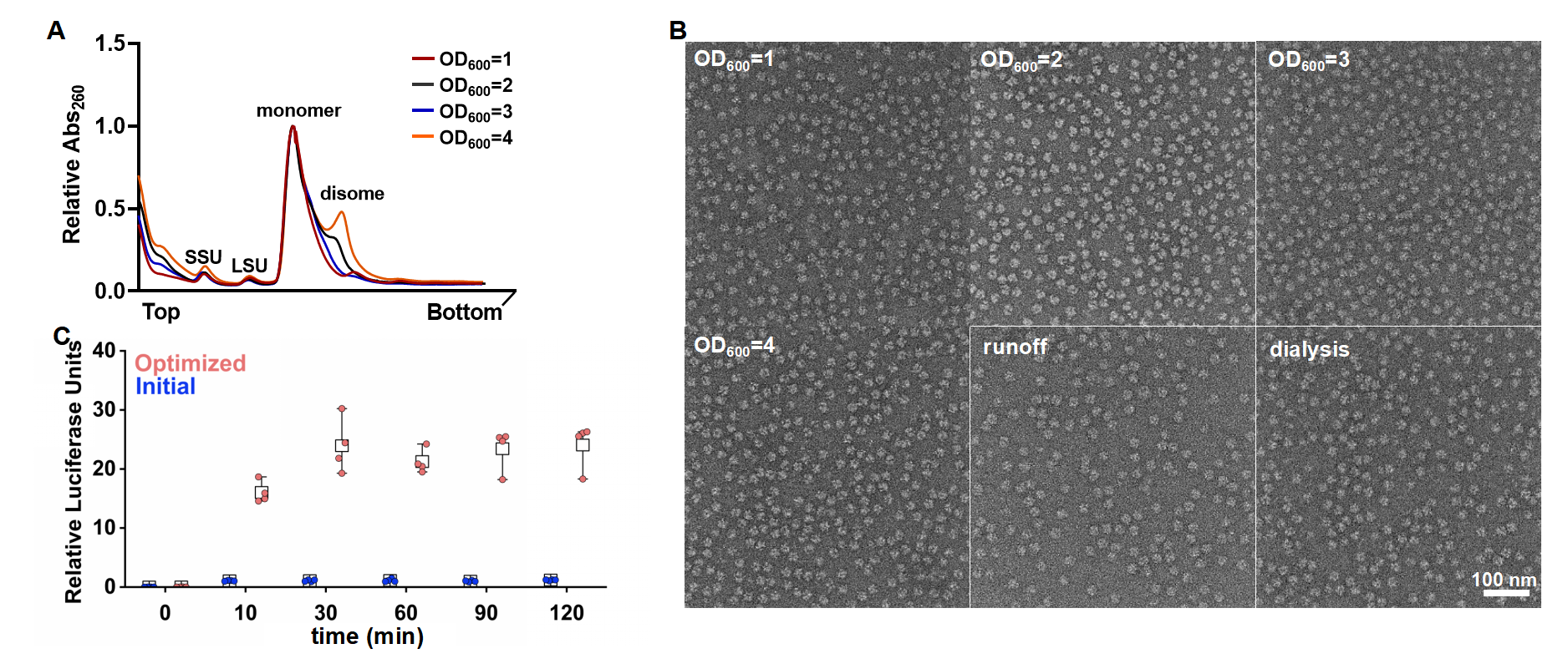

**Figure S5.** **Comparison of different cell lysate preparation methods.** **(A)** Sucrose gradient sedimentation analysis of cell lysates harvested at different cell densities (Optical density, was measured by absorbance at 600 nm). **(B)** Negative-staining TEM images of ribosomes isolated from different lysates. **(C)** Comparison of reaction efficiency in different *e*CFPS systems. Data present mean ± SE, n=4.
